# *Ab initio* predictions for 3D structure and stability of single- and double-stranded DNAs in ion solutions

**DOI:** 10.1101/2022.08.22.504895

**Authors:** Zi-Chun Mu, Ya-Lan Tan, Ben-Gong Zhang, Jie Liu, Ya-Zhou Shi

## Abstract

The three-dimensional (3D) structure and stability of DNA are essential to understand/control their biological functions and aid the development of novel materials. In this work, we present a coarse-grained (CG) model for DNA based on the RNA CG model proposed by us, to predict 3D structures and stability for both dsDNA and ssDNA from the sequence. Combined with a Monte Carlo simulated annealing algorithm and CG force fields involving the sequence-dependent base-pairing/stacking interactions and an implicit electrostatic potential, the present model successfully folds 20 dsDNAs (≤52nt) and 20 ssDNAs (≤74nt) into the corresponding native-like structures just from their sequences, with an overall mean RMSD of 3.4Å from the experimental structures. For DNAs with various lengths and sequences, the present model can make reliable predictions on stability, e.g., for 27 dsDNAs with/without bulge/internal loops and 24 ssDNAs including pseudoknot, the mean deviation of predicted melting temperatures from the corresponding experimental data is only ~2.0℃. Furthermore, the model also quantificationally predicts the effects of monovalent or divalent ions on the structure stability of ssDNAs/dsDNAs.

**Author Summary:** To determine 3D structures and quantify stability of single- (ss) and double-stranded (ds) DNAs is essential to unveil the mechanisms of their functions and to further guide the production and development of novel materials. Although many DNA models have been proposed to reproduce the basic structural, mechanical, or thermodynamic properties of dsDNAs based on the secondary structure information or preset constraints, there are very few models can be used to investigate the ssDNA folding or dsDNA assembly from the sequence. Furthermore, due to the polyanionic nature of DNAs, metal ions (e.g., Na^+^ and Mg^2+^) in solutions can play an essential role in DNA folding and dynamics. Nevertheless, *ab initio* predictions for DNA folding in ion solutions are still an unresolved problem. In this work, we developed a novel coarse-grained model to predict 3D structures and thermodynamic stabilities for both ssDNAs and dsDNAs in monovalent/divalent ion solutions from their sequences. As compared with the extensive experimental data and available existing models, we showed that the present model can successfully fold simple DNAs into their native-like structures, and can also accurately reproduce the effects of sequence and monovalent/divalent ions on structure stability for ssDNAs including pseudoknot and dsDNAs with/without bulge/internal loops.

## Introduction

DNA can adopt many structures beyond the right-handed B-form double-helices, which takes it far beyond being the molecule that stores and transmits genetic information in biological systems (1,2). Some non-B-form DNAs within the human genes, such as hairpins, triplexes, Z-DNA, quadruplexes, and i-motifs, have been proposed to participate in several biologically important processes (e.g., regulation and evolution), leading to mutations, chromosomal translocations, deletions and amplifications in cancer and other human diseases (1–4). Furthermore, self-assembled functional DNA structures have proven to be excellent materials for designing and implementing a variety of nanoscale structures and devices, including interlocked, walkers, tweezers, shuttles, logic circuits, and origami, which have promising applications ranging from photonic devices to drug delivery (5–8). Since short double- and single-stranded DNA (dsDNA and ssDNA) structures (e.g., duplex, hairpins, pseudoknots, and junctions) are essential to build blocks for the construction of non-B-form DNAs and various nano-architectures, advancement in the knowledge of structures and key properties (e.g., thermodynamics and mechanics) for these DNAs will be helpful to understand and ultimately control their biological functions and guide the production and development of novel materials (7–9).

Although several experimental methods such as cryo-electron microscopy, X-ray crystallography, NMR spectroscopy, and other single-molecule techniques (e.g., optical/magnetic tweezers and atomic force microscopy) can be used to determine three-dimensional (3D) structures or elastic properties of DNAs (10–15), there are still full of challenges (e.g., time-consuming and expensive) to experimentally provide insight into DNA folding/hybridization. Thus, the field of computer simulation is rapidly evolving to provide finer details on key features of DNA biophysics compared to experimental approaches (16–19). For example, all-atom molecular dynamics (MD) simulations based on force fields, such as CHARMM and AMBER, have been widely used to investigate dynamics, flexibility, mechanics, or form transition of dsDNA helices at the microscopic level (20–24). However, due to the innumerable degrees of freedom, the MD simulations are limited to small DNAs and to short times even with an advanced-sampling approach and parallel tempering scheme (16,24–26).

On the other hand, the simple continuum DNA models (e.g., worm-like chain model), which treat the double helix as a continuous elastic rod with bending and torsional stiffness, are commonly used to well describe mechanical behavior or elastic bending of dsDNA on long length-scales (27–30). Correspondingly, the nearest-neighbor model can predict secondary structures and melting profiles (e.g., free energy and melting temperature) for ssDNA and dsDNA through the combination of free energy minimization, partition function calculations, and stochastic sampling (9,31). However, these simple models are unable to provide any 3D structure information on DNAs.

Therefore, many coarse-grained (CG) DNA models, which represent DNA using a reduced number of interaction sites while striving to keep the important details, have been developed in recent years to model 3D structures or thermodynamic and structural properties of DNAs (32–39). For example, by mapping each nucleotide into six to seven CG beads, the Martini model combined with MD simulations opens the way to perform large-scale modeling of complex biomolecular systems involving DNA, such as DNA-DNA and DNA-protein interactions (40,41). Very recently, a three-bead CG model, MADna, was developed to reproduce the conformational and elastic features of dsDNA, including the persistence length, stretching/torsion moduli, and twist−stretch couplings (42). However, since the two models need constraints (e.g., predefined elastic network or bonded interactions for paired bases) to maintain a double helix, they cannot be used to study DNA hybridization, melting, and hairpin formation (40–42).

Moreover, some other Go-like models, including 3SPN, oxDNA, and TIS, have been proposed to fill the gap (43–50). The 3SPN model, which reduces the complexity of a nucleotide to three interactions sites (i.e., phosphate, sugar, and base), can successfully capture DNA denaturation/renaturation and provide a reasonable description of other thermomechanical and structural properties for DNAs (e.g., persistence length, bubble formation, major and minor groove widths, and local curvature) by involving in base-stacking and base-pairing interactions (43–45). The oxDNA model uses three collinear sites and a vector normal to the base site to construct the angle-dependent potentials including coplanar base-stacking and linear hydrogen bonding interactions, which are parametrized to accurately describe basic structural, mechanical, and thermodynamic properties of ss/dsDNA (46–48). More significantly, with fine-tuned structural parameters, the model can also treat large DNA nanostructures, such as DNA origami and nanotweezers (48,49). The TIS-DNA is another robust three-interaction-site CG model, and using a set of nucleotide-specific stacking parameters obtained from thermodynamic properties of dimers, the model can reproduce the sequence-dependent mechanical, as well as thermodynamic properties of DNAs, covering the force-extension behavior and persistence lengths of poly(dA)/poly(dT) chains, elasticity of dsDNA and melting temperatures of hairpins (50–52). The use of Go-like interactions (e.g., non-bonded potentials to penalize deviations from a reference structure) can effectively restrict the range of conformations that may be sampled by the CG model, and simultaneously, it also limits the possibility of the model on structure prediction from the sequence.

Recently, the 3dRNA/DNA web server was further developed based on the 3dRNA to build three-dimensional (3D) structures of RNA and DNA from template segments with very high accuracy using sequence and secondary structure information (53,54). Similarly, a pipeline presented by Jeddi and Saiz can also be used to predict DNA hairpins by integrating the existed 2D and 3D structural tools (e.g., Mfold, Assemble, and MD) (55). However, the two structure prediction methods are dependent on secondary structures, while there is still a problem to exactly predict secondary structures of DNAs (31). Fortunately, a minimal physics-based CG model of nucleic acids named NARES-2P was proposed to fold dsDNA from separate strands without any Go-like potentials and secondary structure information. Although the model was constructed using the bottom-up strategy, where each component of the energy function was fitted separately to the respective potential of mean force obtained from all-atom potential-energy surfaces, it can reproduce many properties of double-helix B-DNA, such as duplex formation, melting temperatures, and mechanical stability (56–58). Contrary to the oversimplification of NARES-2P (i.e., two sites per nucleotide), the HiRE-RNA is an empirical CG model for RNA and DNA, whose resolution is high enough (i.e., six or seven beads for each nucleotide) to preserve many important geometric details, such as base pairing and stacking. Without imposing preset pairings for the nucleotides, the HiRE-RNA can investigate both dynamical and thermodynamic aspects of dsDNA assemblies, as well as the effect of sequences on the melting curves of the duplexes (59,60). Despite the advances, the parameters of the two models may need further validation for quantifying thermodynamic and 3D structure to accord with experiments, especially for ssDNA.

In addition, due to the polyanionic nature of DNAs, metal ions (e.g., Na^+^ and Mg^2+^) in solutions can play an essential role in DNA folding and dynamics (12–14,61–64). Although several of the existing models such as 3SPN, oxDNA, TIS, and NARES-2P have taken the electrostatic interactions into account using the Debye-Huckel approximation or mean-field multipole–multipole potentials and reproduced some monovalent salt-dependent structural properties (e.g., persistence length, torsional stiffness or melting temperatures) of DNAs (45,48,50,57), quantitatively predicting the 3D structure and thermodynamic stability for DNA including both ssDNA and dsDNA in ion solutions (especially divalent ions) from the sequence is still an unresolved problem. Recently, we have proposed a three-bead CG model to simulate RNA folding from the sequence, and with an implicit electrostatic potential, the model can make reliable predictions on 3D structures and stability for RNA hairpins, pseudoknots, and kissing complexes in ion solutions (65–69). However, due to the differences in geometry, base stacking strength, and flexibility between DNA and RNA, the present model cannot be directly used to simulate DNA folding.

In this work, we further developed an *ab initio* CG model of DNA to predict the 3D structure, stability, and salt effect for both dsDNA and ssDNA. First, the bonded and nonbonded potentials were parameterized based on the statistical analysis of known DNA 3D structures, as well as experimental thermodynamic parameters and melting data. Afterward, the model was validated through 3D structure and stability predictions for DNAs including double helices, hairpins, and pseudoknots with different lengths and sequences, as compared with the extensive experimental data. Furthermore, we also showed that the effects of monovalent and divalent ions on DNA structure stability predicted by the present model are in accordance with the corresponding experiments.

## Materials and methods

### CG structure representation for DNAs

To be consistent with our previous RNA CG model (65), each nucleotide in DNA is also simplified into three beads: P, C, and N, to represent the phosphate group, sugar ring, and base plane, respectively. For simplicity, the three beads are placed at the positions of existing atoms (i.e., P, C4’, and N1 for pyrimidine or N9 for purine) (Fig. 1) and are treated as van der Waals spheres with the radii of 1.9Å, 1.7Å and 2.2 Å, respectively (65,69). One unit negative charge (-*e*) is placed on the center of P bead to describe the highly charged nature of DNA.

**Figure 1.**
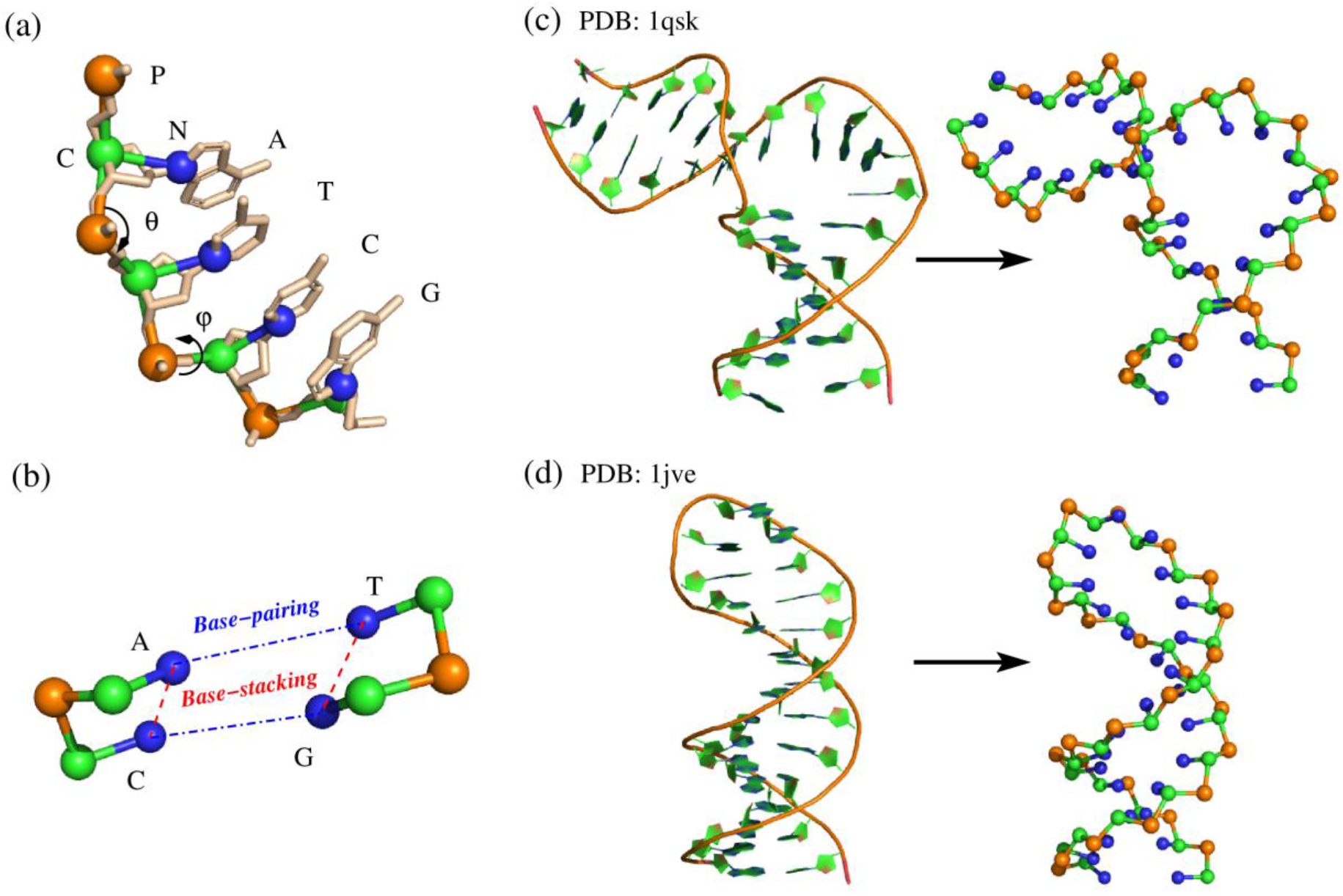
The representations of all-atom and CG model for each deoxynucleotide and ss/dsDNA molecules, as well as the schematics of base-pairing and base-stacking in the present model. (a) Our coarse-grained representation of a DNA fragments including deoxynucleotides of A, T, G, and C superimposed on the all-atom representation. The three beads are located at phosphate (P), sugar (C4′) and pyrimidine (N1) or purine (N9). θ and φ are the schematics of CG bonded angle (CPC) and dihedral (CPCP), respectively. (b) The schematic representation of base-pairing (blue) and base-stacking (red) interaction. (c, d) The 3D structures of (c) a dsDNA with bulge loop (PDB:1qsk) and (d) an ssDNA hairpin (PDB: 1jve) in all-atomistic (left) and our CG representation (right). The 3D structures are shown with PyMol (http://www.pymol.org).

### Energy functions

The total energy *U* in the present DNA CG model is composed of the following eight components:

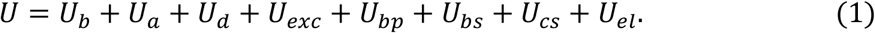

The first three terms are bonded potentials for virtual bonds *U*_*b*_, bond angles *U*_*a*,_ and dihedrals *U*_*d*_, respectively, which are used to mimic the connectivity and local geometry of DNA chains. The function forms of these terms are listed in the Supporting information, which can also be found elsewhere (44,46,50,65).

The remaining terms of Eq. 1 describe various pairwise, nonbonded interactions. The *U*_*exc*_ represents the excluded volume interaction between the CG beads and it is modeled by a purely repulsive Lennard-Jones potential. The *U*_*bp*_ in Eq. 1 is an orientation-dependent base-pairing interaction for the possible canonical Watson-Crick base pairs (G-C and A-T). The formula of *U*_*bp*_ is similar to the form of hydrogen-bonding interaction used in the TIS model (50), and the backbone dihedrals are replaced by two simpler distances between CG beads in pairing nucleotides to describe the orientation of hydrogen-bonding interactions; see Eq. S6 in the Supporting information. The sequence-dependence base-stacking interaction *U*_*bs*_ between two nearest neighbour base pairs is given by

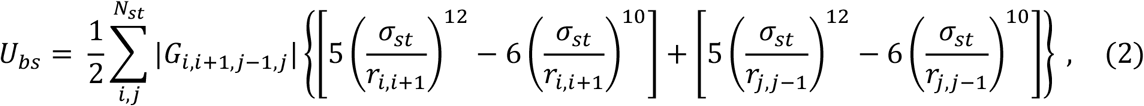

where *σ*_*st*_ is the optimum distance of two neighbour bases in the known DNA helix structures. *G*_*i*,*i*+1,*j*−1,*j*_ in Eq. 2 is the strength of base-stacking energy, and it can be calculated by *G*_*i*,*i*+1,*j*−1,*j*_ = Δ*H* – *T*(Δ*S* – Δ*S*_*c*_). Here, *T* is the absolute temperature in Kelvin, Δ*H* and Δ*S* are the DNA thermodynamic parameters derived from experiments (9,70), and Δ*S*_*c*_ is the conformational entropy change that is naturally included in the Monte Carlo (MC) simulations, during the formation of one base pair stacking; see more details in Eq. S7 and Fig. S3 in the Supporting information as well as our previous works (65,69). In addition, the coaxial-stacking interaction *U*_*cs*_ between two discontinuous neighbour helices is also taken into account by the present model through calculating the stacking potential of the interfaced base-pairs, and the expression can be found in Eq. S10 in the Supporting information.

The last term *U*_*el*_ in Eq. 1 is used to calculate the electrostatic interactions between phosphates groups (e.g., *i*-th and *j*-th P beads), and it is given by

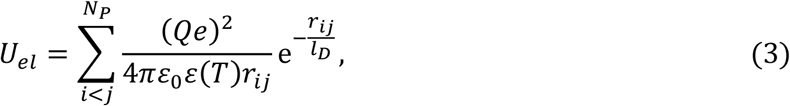

where *e* is the elementary charge, *r*_*ij*_ is the distance between *i*- and *j*-th P beads, and *N*_P_ is the total number of P beads in a DNA. *l*_*D*_ is Debye length, which defines the ionic screening at different solution ionic strengths. *ε*_0_ and *ε*(*T*) are the permittivity of vacuum and an effective temperature-dependent dielectric constant, respectively (50,65,66). *Q* is the reduced charge fraction derived based on Manning’s counterion condensation theory and the tightly bound ion model (71–73); see Eq. S11 in the Supporting information. Due to the inclusion of the *U*_*el*_, the present model can be used to study DNA folding in pure (e.g., Na^+^) as well as mixed (e.g., Na^+^/Mg^2+^) ion solutions.

### Parametrization

The initial parameters of bonded potentials (i.e., *U*_*b*_, *U*_*a*,_ and *U*_*d*_ in Eq. 1) were derived from the Boltzmann inversion of the corresponding CG atomistic distribution functions, obtained by the statistical analysis on experimental DNA structures in the Protein Data Bank (PDB) (http://www.rcsb.org/pdb/-home/home.do) (Fig. S1), and the PDB codes of 138 DNA structures used in this work are listed in Table S1 in the Supporting information. Note that these DNAs are with no overlap with ssDNAs/dsDNAs used for model validation on 3D structure prediction. Since the known DNA structures are generally double helices, the initial parameters from these structures could not be reasonable to describe DNA chains during folding processes. In our previous RNA model, two sets of parameters (Para_helical_ and Para_nonhelical_) were calculated from stems and loops in experimental structures, respectively (65,66), and the Para_nonhelical_ ones were used to successfully describe the folding of an RNA from a free chain. However, due to the limitation of the number of loop regions in known DNA structures, obtaining suitable parameters for DNA-free chains directly from these structures is unrealistic. Although we also did MD simulations for unstructured ssDNA (e.g., polyA and polyT chains) and tried to extract the bonded parameters from the conformations (data not shown here), because of some differences in optimum values of several angles between experimental and MD simulated structures, we gave them up. Eventually, based on the distributions of bond length/angle/dihedral for nonhelical parts in RNA structures are just slightly broader than that of helical parts (65), we simply set the strengths of DNA bonded potentials in Para_nonhelical_ as one-half of that in Para_helical_. Note that, the Para_nonhelical_ is used in the folding process, and the Para_helical_ is only used for stems during folded structure refinement. Whereafter, the initial parameters were further optimized through the comparisons between the simulated and experimental bond length/angle distributions (34,74), and in this process, there are only two dsDNAs (PDB codes: 1agh, 3bse) and two ssDNAs (PDB codes: 1ac7, 1jve) were used.

For nonbonded potentials, the geometric parameters in base-pairing/stacking functions were obtained from the known structures; see Fig. S2 in the Supporting information. The strength of base-stacking was estimated from the combination of the experimental thermodynamics parameters and MC simulations; see Eqs. S7-S9 and Fig. S3 in the Supporting information. The strength of base-pairing (i.e., *ε*_*bp*_ in Eq. S6) was determined by comparing the predicted melting temperatures (*T*_m_’s) of four ss-/dsDNAs with corresponding experiments. That is, for two ssDNA hairpins (sequences: GCGCTTTTTGCGC and GGAGCTTTTTGCTGC; ion condition: 1M NaCl; see Table 1) and two dsDNAs (sequences: GCTAGC/GCTAGC and GGGACC/GGTCCC; strand concentration: 0.1mM and 0.4mM, respectively; ion condition: 1M NaCl; see Table 2), we used the present model to predict their *T*_m_’s through continuously adjusting the *ε*_*bp*_ until the agreement between predicted and experimental data is satisfactory.

**Table 1.**
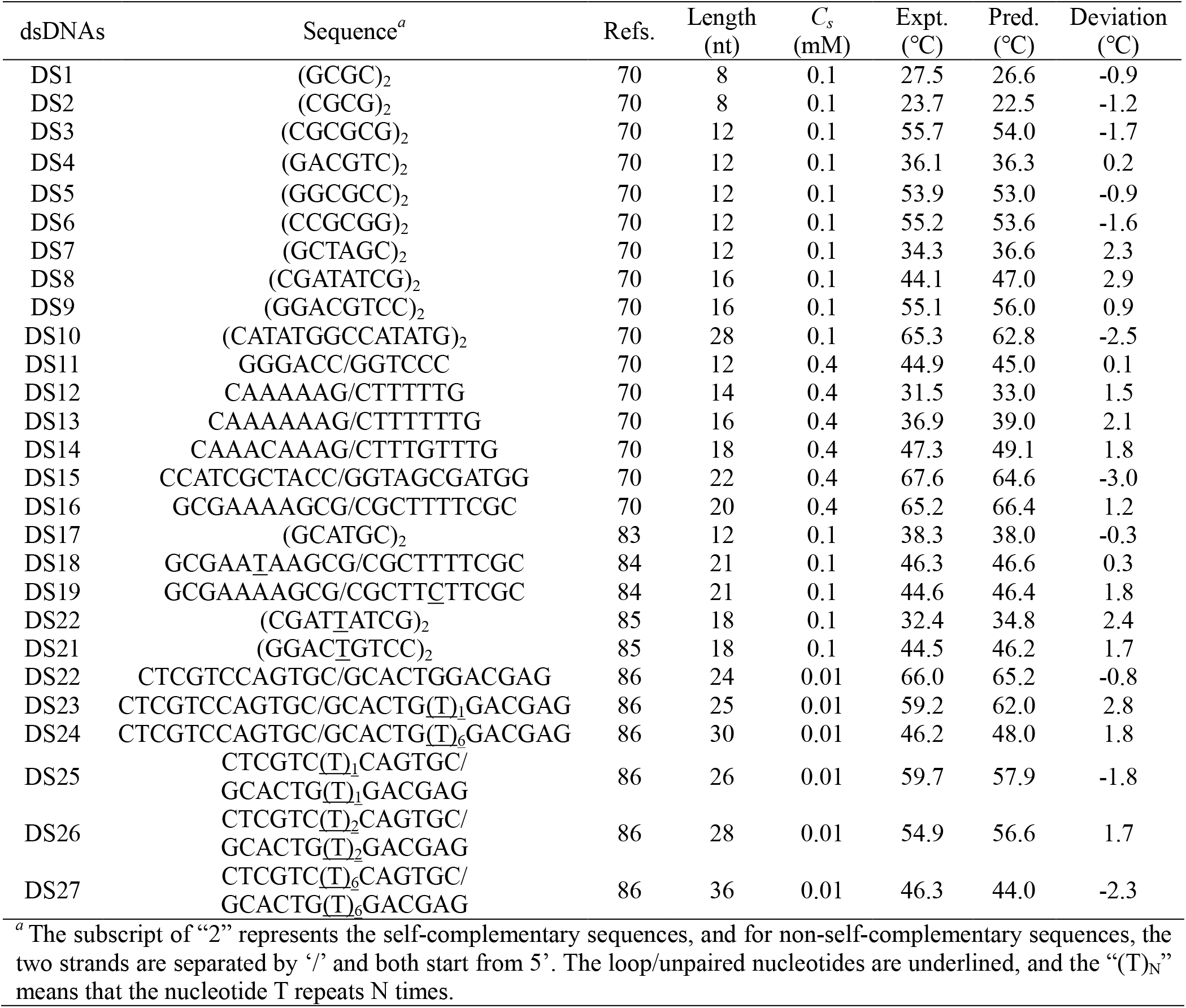
The melting temperatures (*T*_m_) for dsDNAs at 1M [Na^+^].

**Table 2.** The melting temperatures (*T*_m_) for single-stranded DNAs at given ion conditions

| ssDNAs | Sequence<sup>a</sup> | Refs. | Length<br>(nt) | [Na<sup>+</sup>]<br>(M) | Expt.<br>(°C) | Pred.<br>(°C) | Deviation<br>(°C) |
| --- | --- | --- | --- | --- | --- | --- | --- |
| SS1 | GCGC(T)<sub>5</sub>GCGC | 88 | 13 | 1.0 | 76.4 | 78.4 | 2.0 |
| SS2 | GCCGC(T)<sub>5</sub>GCGGC | 88 | 15 | 1.0 | 81.6 | 84.5 | 2.9 |
| SS3 | GCAGC(T)<sub>5</sub>GCTGC | 88 | 15 | 1.0 | 74.9 | 75.8 | 0.9 |
| SS4 | GCTGC(T)<sub>5</sub>GCGC | 88 | 15 | 1.0 | 58.0 | 59.0 | 1.0 |
| SS5 | GCTGC(T)<sub>5</sub>GCTGC | 88 | 15 | 1.0 | 56.9 | 55.1 | -1.8 |
| SS6 | GCGC(T)<sub>3</sub>GCGC | 87 | 11 | 0.1 | 80.7 | 78.0 | -2.7 |
| SS7 | GCGC(T)<sub>5</sub>GCGC | 87 | 13 | 0.1 | 73.3 | 71.9 | -1.4 |
| SS8 | GCGC(T)<sub>7</sub>GCGC | 87 | 15 | 0.1 | 62.8 | 63.9 | 1.1 |
| SS9 | GCGC(T)<sub>9</sub>GCGC | 87 | 17 | 0.1 | 56.2 | 58.1 | 1.9 |
| SS10 | GTAC(T)<sub>5</sub>GTAC | 89 | 13 | 0.1 | 33.4 | 32.2 | -1.2 |
| SS11 | ATCCTA(T)<sub>5</sub>TAGGAT | 90 | 17 | 0.2 | 51.0 | 54.0 | 3.0 |
| SS12 | ATCCTA(T)<sub>6</sub>TAGGAT | 90 | 18 | 0.2 | 48.1 | 51.6 | 3.5 |
| SS13 | ATCCTA(T)<sub>7</sub>TAGGAT | 90 | 19 | 0.2 | 43.9 | 47.1 | 3.2 |
| SS14 | GAATTC(T)<sub>5</sub>GAATTC | 91 | 17 | 0.2 | 55.3 | 53.1 | -2.2 |
| SS15 | CCCAA (T)<sub>12</sub>TTGGG | 92 | 22 | 0.25 | 57.9 | 59.2 | 1.1 |
| SS16 | CCCAA (T)<sub>16</sub>TTGGG | 92 | 26 | 0.25 | 51.7 | 55.5 | 3.8 |
| SS17 | CGGATAA(T)<sub>4</sub>TTATCCG | 93 | 18 | 0.1 | 63.4 | 66.0 | 2.6 |
| SS18 | CGGATAA(T)<sub>8</sub>TTATCCG | 93 | 22 | 0.1 | 50.5 | 52.5 | 2.0 |
| SS19 | CGGATAA(T)<sub>12</sub>TTATCCG | 93 | 26 | 0.1 | 41.1 | 42.0 | 0.9 |
| SS20 | CGGATAA(T)<sub>16</sub>TTATCCG | 93 | 30 | 0.1 | 35.1 | 34.1 | -1.0 |
| SS21 | CGGATAA(T)<sub>20</sub>TTATCCG | 93 | 34 | 0.1 | 29.6 | 26.8 | -2.8 |
| SS22 | AAAAAAA(C)<sub>5</sub>TTTTTTT | 36 | 19 | 1.0 | 55.0 | 58.5 | 3.5 |
| SS23 | CGCGCG(T)<sub>4</sub>GAAATTCGCGCG | 87 | 32 | 0.1 | 52.6/ | 48.8/ | -3.8/ |
|  | (T)<sub>6</sub>AATTTC |  |  |  | 70.7 | 72.0 | 1.3 |
| SS24 | GCGC(T)<sub>5</sub>GCGCGTAC(T)<sub>5</sub>GTAC | 89 | 26 | 0.1 | 50.1/ | 46.9/ | -2.1/ |
|  |  |  |  |  | 76.1 | 75.0 | -1.1 |

<sup>a</sup> The sequences start from 5', the loop nucleotides are underlined, and the "(T)<sub>N</sub>" means that the nucleotide T repeats N times.

The detailed descriptions, as well as the parameters of all the potentials in Eq. 1, can be found in the Supporting information.

### Simulation procedure

During DNA folding from sequence without any preset constraints, it is easy to fall into a metastable state with local minimum energy. To effectively avoid that, the MC simulated annealing algorithm, whose capacity has been proved in protein/RNA folding, was used to sample conformations for ssDNA or dsDNA (65,75,76). For each DNA, a random chain configuration is generated from its sequence, and for dsDNA, the two chains are separately placed in a cubic box, the size of which is determined by the concentration of a single strand. Afterward, the simulation of a DNA system with a given monovalent/divalent ion condition is performed from a high temperature (e.g., 120℃) to the target temperature (e.g., room/body temperature). At each temperature, conformational changes are accomplished via the translation and pivot moves, which have been demonstrated to be rather efficient in sampling conformations of polymers (77,78), and the changes are accepted or rejected according to the standard Metropolis algorithm (65,69). The equilibrium conformations at different temperatures during the cooling process are used to analyze the stability of the DNA. In structure prediction, the last conformation at the target temperature is taken as the initial predicted structure, which can be further refined to better capture the geometry of helical parts by introducing the bonded parameters of Para_helical_ for consecutive base-pairing regions. After structure refinement, an ensemble of structures would be obtained, and the mean RMSD (the averaged value over the whole structure ensemble) and minimal RMSD (corresponding to the structure closest to the native one) calculated over CG beads from the corresponding atoms in the native PDB structure is used to evaluate the reliability of the present model on DNA 3D structure prediction.

### Calculation of melting temperature

At each temperature, the fractions of folded (F, consistent secondary structures with predicted at lowest temperature) and unfolded (U, no more than one base pair) states could be fitted to a two-state model through the following equations (9,65):

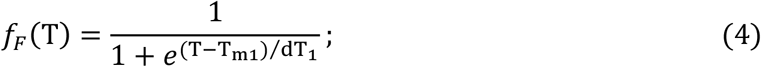

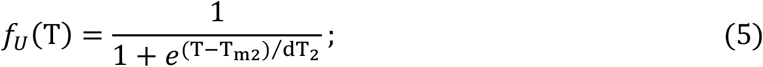

where *T*_m1_ and *T*_m2_ are the two melting temperatures of the corresponding transitions (folded state to possible intermediate state (I) and intermediate state to unfolded state), respectively. *dT*_1_ and *dT*_2_ are the corresponding adjustable parameters. Based on the *f*_F_(T) and *f*_U_(T), the fraction of the number of denatured base pairs *f*(T) can be calculated by (68)

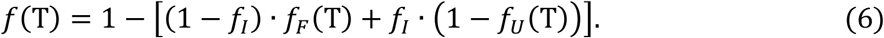

Here, *f*_*I*_ is the fraction of the number of denatured base pairs when the fraction for the I state is maximum. And then, the d*f/*d*T* (the first derivative of *f* with respect to temperature) profile can be calculated to compare with the corresponding experimental data. It should be noted that for simple hairpins and short duplexes used in this work, the I state almost never occurs, and the *f*_*I*_ in Eq. 6 could be set to 0, which means that *f*_*U*_(T) is approximately equal to 1 – *f*_*F*_(T) and only one *T*_m_ can be obtained.

To improve the simulation efficiency for dsDNA with low strand concentrations *c*_*s*_ (e.g., <0.1mM), the MC simulations were performed at a relatively high strand concentration *c*_*s*_^*h*^ (e.g., 1mM), and the fraction (*f*(T; *c*_*s*_)) of denatured base pairs at lower *c*_*s*_ can be calculated by that (*f*(T; *c*_*s*_^*h*^)) at *c*_*s*_^*h*^ (69,79,80)

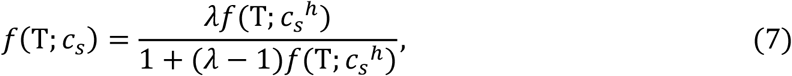

where *λ* = *c*_*s*_^*h*^/*c*_*s*_. Furthermore, for a dsDNA with a two-state transition, the melting temperature *T*_*m*_(*c*_*s*_) at *c*_*s*_ can be directly obtained from *T*_*m*_(*c*_*s*_^*h*^) based on Eqs. 4–7,

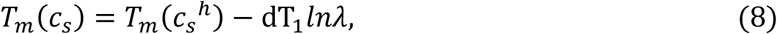

the derivation of which can be found in the Supporting information.

## Results

Based on the parameterized implicit-solvent/salt energy function and the MC simulated annealing algorithm, the present CG model can be used to predict 3D structures for dsDNA as well as ssDNA at different ion conditions and temperatures from the sequence. In this section, we tested the present model on the 3D structure and stability predictions for extensive DNAs with various lengths/sequences. As compared with the experimental structures and thermodynamics data, the present model can make overall reliable predictions.

### DNA 3D structure prediction from sequence

#### For dsDNAs

As described in the section of “Material and methods”, for each dsDNA, two random chains were generated from its sequence (e.g., structure A in Fig. 2a), which were further randomly placed in a cubic box, ensuring that there is no overlap. To guarantee no significant effect of the box on 3D structures, the strand concentration was set as 1mM (i.e., box side length of 149 Å) for short dsDNA (<10bp) and 0.1mM for longer ones. Due to the lack of the ion conditions for the experimental structures determined by X-ray crystallography, for simplicity, we only predicted the 3D structures for all DNAs at high ion concentrations (e.g., 1M NaCl), regardless of possible ion effects. As shown in Fig. 2a, for a dsDNA with a five-adenine bulge loop (PDB: 1qsk; 29nt, 12bp), the energy of the system reduces with the formation of base pairs as the temperature is gradually decreased from 120℃ to 25℃, and the initial random chain folds into its native-like double-stranded structures (e.g., structure C in Fig. 2a). Following that, another MC simulation (e.g., 1×10^5^ steps) is performed at target temperature based on the final structure predicted by the preceding annealing process, and the two sets of bonded potential parameters Para_nonhelical_ and Para_helical_ are employed respectively for the single-strands/loops and base-pairing regions to better capture the geometry of the helical part. As shown in the inset of the bottom panel of Fig. 2a, the mean and minimum RMSDs of the dsDNA between predicted structures and its native structure are ~3.2Å and ~1.8Å, respectively, and the corresponding predicted 3D structures are as shown in Fig. 3a.

**Figure 2.**
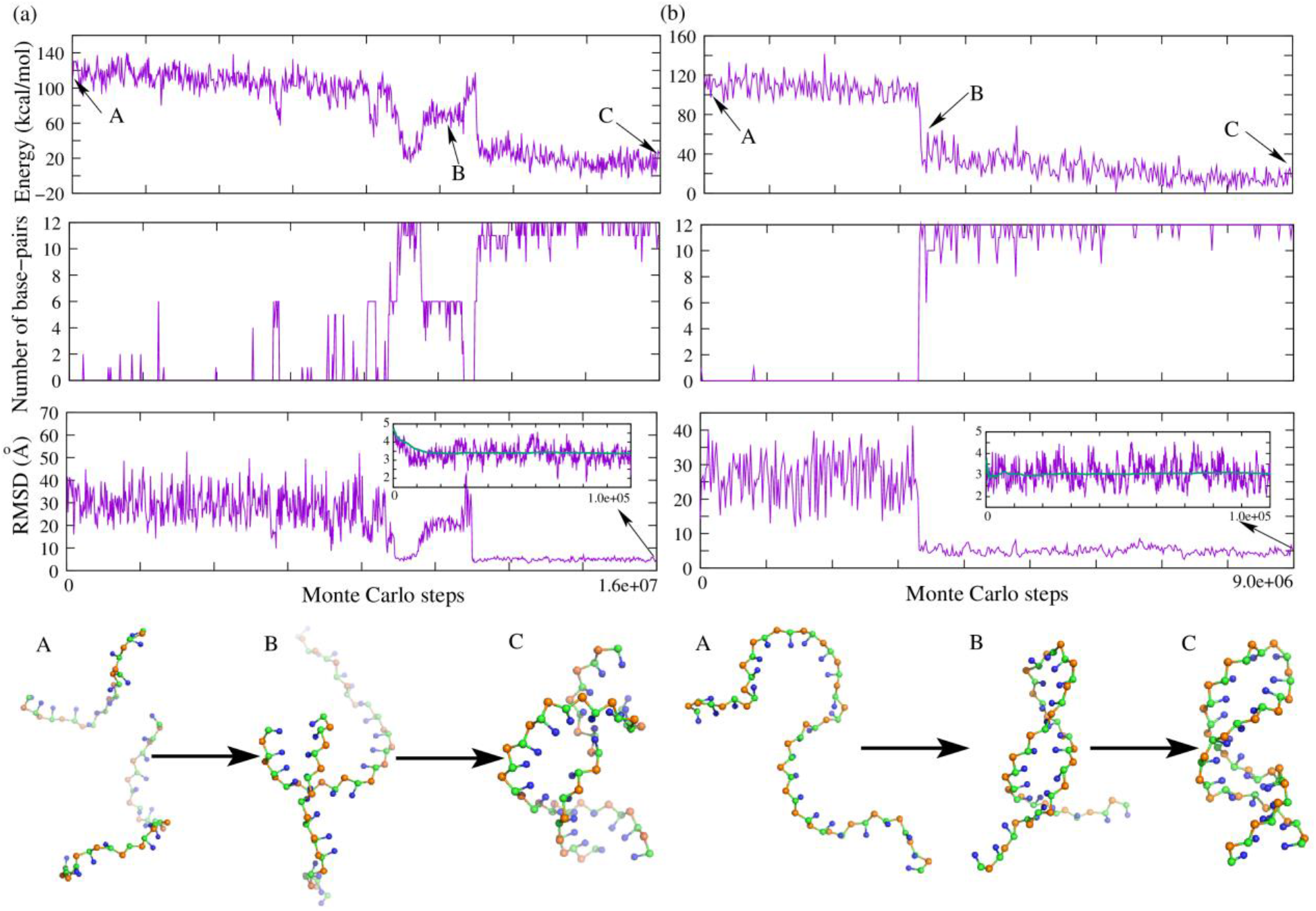
3D structure prediction for the paradigm dsDNA/ssDNA in the present model. (a, b) The time-evolution of system energy, number of base-pairs, RMSD from native structure, and typical 3D conformations (from top to bottom, respectively) during the Monte Carlo simulated annealing simulation for (a) a dsDNA (PDB: 1qsk) and (b) an ssDNA (PDB: 1jve). The insets show the RMSDs of refined conformations calculated over all CG beads from the corresponding atoms in native structures. The 3D structures are shown with PyMol (http://www.pymol.org).

**Figure 3.**
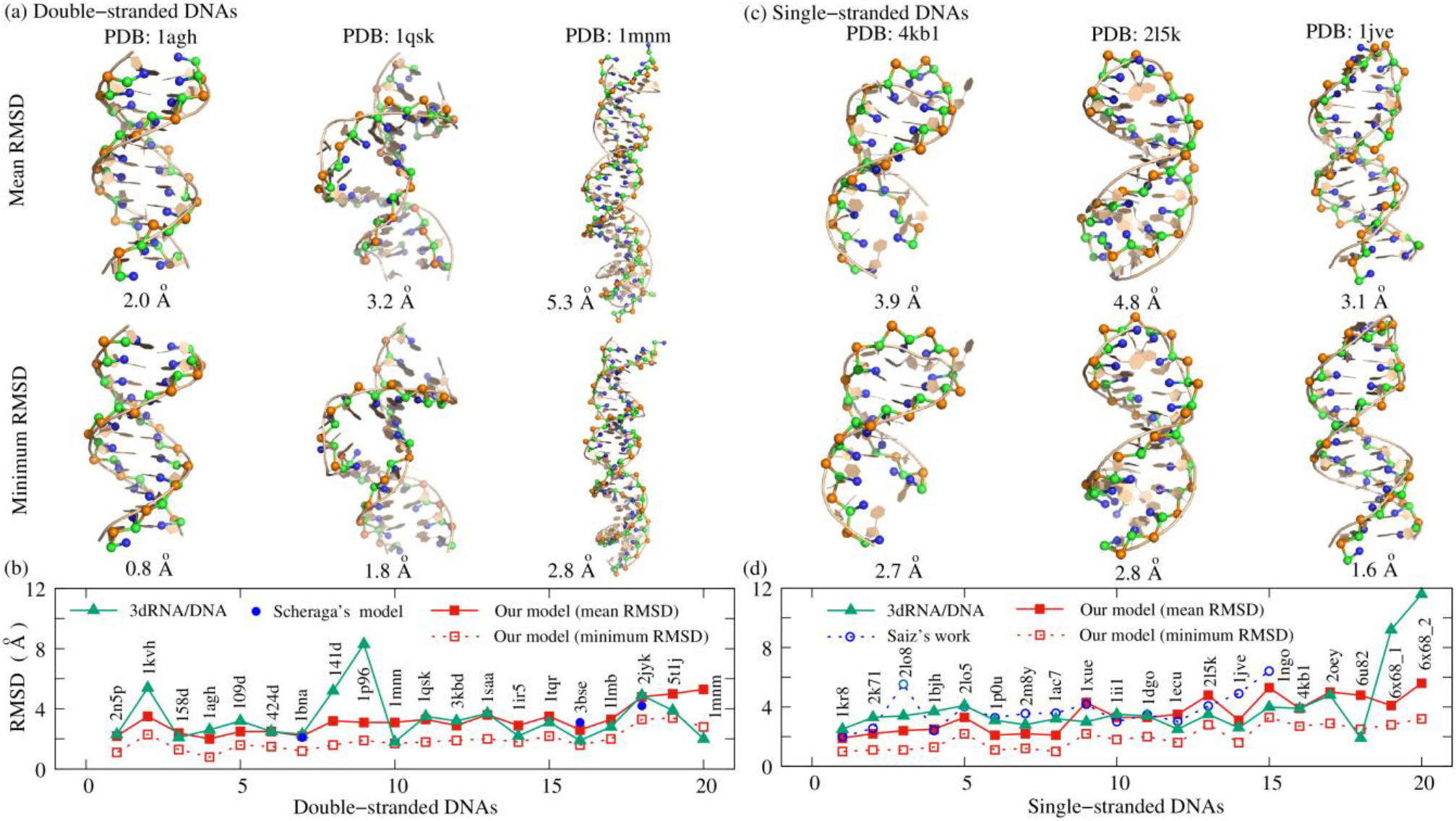
The display of typical predicted 3D structures, and comparisons of RMSDs between the present model and other models. (a, c) The predicted 3D structures (ball-stick) with the mean (top) and minimum (bottom) RMSDs for (a) three sample dsDNAs (PDBs: 1agh, 1qsk, and 1mnm) and (c) three ssDNAs (PDBs: 4kbl, 2l5k, and 1jve) from their native structures (cartoon). (b, d) The comparison of the predicted structures between the present model and the existing models including the 3dRNA/DNA and the models from Scheraga’s or Saiz’s group for (b) 20 dsDNAs and (d) 20 ssDNAs. The results of 3dRNA/DNA are predicted through their online server based on the native secondary structures (53). The other data is taken from refs. 55 and 58. The 3D structures are shown with PyMol (http://www.pymol.org).

According to the above process, we employed the present model to predict the 3D structures of 20 dsDNAs (18nt-52nt) including helix with bulge loops, and the detailed descriptions (e.g., sequence, length, and structure feature) of these dsDNAs are listed in Table S3 in the Supporting information. For the 20 dsDNAs, the overall mean and minimum RMSD values are ~3.2 Å and ~1.9 Å, respectively; see Fig. 3a and Table S3 in the Supporting information, which suggest that the present model can make reliable predictions for 3D structures of dsDNA just from the sequence, despite a certain deviation (especially at the two ends) between the predicted and experimental structures for large dsDNA (e.g., PDB: 1mnm and 5t1j). Fig. 3a also shows the predicted 3D structures (ball-stick) with the mean and minimum RMSDs and the experimental structures (cartoon) for three typical dsDNAs with different lengths and sequences, intuitively indicating the ability of the model.

#### For ssDNAs

Compared with most of the existing models, the present model cannot only predict the double helix structure of dsDNA, but it can also make a prediction on the 3D structure for more flexible ssDNA. Similarly, a random chain generated from one ssDNA sequence can fold into native-like structures with temperature dropping; see Fig. 2b for an example of a DNA hairpin (PDB: 1jve; 27nt, 12bp), which could primarily benefit from the use of the soft parameters (Para_nonhelical_) of bonded potentials and sequence-dependent base-stacking interactions in the present model. As shown in Fig. 3b, for 20 ssDNAs (7nt-74nt) used in this work including hairpins with bulge/internal loops (Table S4 in the Supporting information), the overall mean (minimum) RMSD between the predicted and experimental structures is ~3.5Å (~2.0Å), which strongly suggested that the present model can successfully predict 3D structures for simple ssDNA.

Since the structures of the largest hairpin (i.e., 6×68_2) from the *piggyBac* DNA transposon (PDB: 6×68, a synaptic protein-DNA complex) has a significant bending possibly influenced by protein (81), our predictions without regard to protein has a certain deviation (mean RMSDs of 5.6Å) from the experimental structure; see Fig. 3d. It is worth noting that beyond DNA hairpins, we also tried to predict the 3D structure for a DNA three-way junction using the present model. As shown in Fig. S4 in the Supporting information, the structures (two hairpins at two ends) predicted from the sequence are pretty inconsistent with experimental ones. To find why, we further performed a MC simulation using the present model for the ssDNA starting from its PDB structure, and found that there is no significant difference in energies between predicted and simulated conformations (Fig. S4), which suggests that some tertiary interactions including the noncanonical base-pairing and base-backbone hydrogen bonding (82) and a more efficient algorithm (e.g., replica-exchanged MC) should be further taken into account in the model.

#### Comparisons with other models

To further examine the ability of the model on predicting 3D structures of DNAs (ssDNA and dsDNA), we also made comparisons with available results from the existing models. First, we employed the 3dRNA/DNA web server (http://biophy.hust.edu.cn/new/3dRNA/create), which is an automatic, fast, and high-accuracy RNA and DNA tertiary structure prediction method (53,54), to predict 3D structures for all DNAs used in this work using the default options (e.g., Procedure: best; Loop Building Method: Bi-residue; # of Predictions: 5) and experimental secondary structures, and calculated the mean RMSD of returned conformations for each DNA over the atoms of P, C4’ and N1/N9 from the corresponding atoms in the experimental structures. As shown in Fig. 3, for 20 dsDNAs, the overall mean RMSD (~3.2 Å) from the present model is not worse than that (~3.3 Å) from the 3dRNA/DNA, and for 20 ssDNAs, our prediction (overall mean RMSD: ~3.5 Å) is slightly smaller than predicted result (~4.0 Å) from the 3dRNA/DNA.

Furthermore, we also made comparisons with the predictions from Refs. 55 and 58. Scheraga et al. also proposed a physics-based rigid-body CG model (3-bead) of DNA, and used it to successfully fold 3 dsDNAs (PDBs: 1bna, 3bse, and 2jyk) from complementary strands with only weak constraints between them (58). The all-bead RMSDs of the three lowest-energy predicted structures with respect to experimental references are 2.1Å, 3.1Å, and 4.2Å, respectively, which are close to the mean RMSDs (2.2Å, 2.6Å, and 4.8Å, respectively) predicted from the present model (Fig. 3a). Jeddi and Saiz presented a pipeline that integrates sequentially building ssDNA secondary structure from Mfold, constructing equivalent 3D ssRNA models by Assemble 2, transforming the 3D ssRNA models into ssDNA 3D structures, and refining the resulting ssDNA 3D structures through MD simulations (55). As shown in Fig. 3b, for 15 ssDNA hairpins, the average RMSD (over the sugar-phosphate backbone) for the best structures predicted by the pipeline is ~3.7Å, a visibly larger value than the overall mean/minimum RMSD (~3.2Å/~2.2Å) from our predictions. Therefore, the comparisons with the other models fully show that the present model can successfully fold simple dsDNA/ssDNA from the sequence without the help of any secondary structure information.

### Stability of various DNAs

Beyond 3D structure predictions, the present model can also be used to predict the thermal stability for dsDNA and ssDNA in ion solutions. In order to verify the effect of the model, we further used it to predict the melt temperatures for extensive DNAs.

#### For dsDNA with various lengths/sequences

The melting temperature (*T*_m_) of each dsDNA can be calculated by the present model based on 3D structures predicted at different temperatures; see Fig. 4a and the section “Material and methods”. For example, for the sequence (GGACGTCC)_2_ at 1M [Na^+^], the melting curve of the dsDNA with a high strand concentration of 1 mM was predicted according to the fractions of unfolded state at different temperatures (Eqs. 5–7), and the melting curve, as well as the *T*_m_ of the dsDNA at low experimental strand concentration (0.1 mM), can be obtained through Eq. 8; see Figs. 4a and b. The predicted *T*_m_ of the sample sequence at *c*_*s*_ = 0.1mM is ~56.0°C, which is only 0.9°C higher than the corresponding experimental value (~55.1°C) (70). We further performed simulations for the dsDNA at *c*_*s*_ = 0.1mM to directly predict its *T*_m_ at experimental strand concentration, and found that there is no significant difference between two melting curves/temperatures (Fig. 4b). In addition, as shown in Fig. 4c, the predicted *T*_m_’s for three different dsDNAs at different strand concentrations are also in good accordance with the experiments (70), proving that it is feasible to infer the *T*_m_ at low *c*_*s*_ from the high ones (*c*_*s*_^*h*^) (79,80).

**Figure 4.**
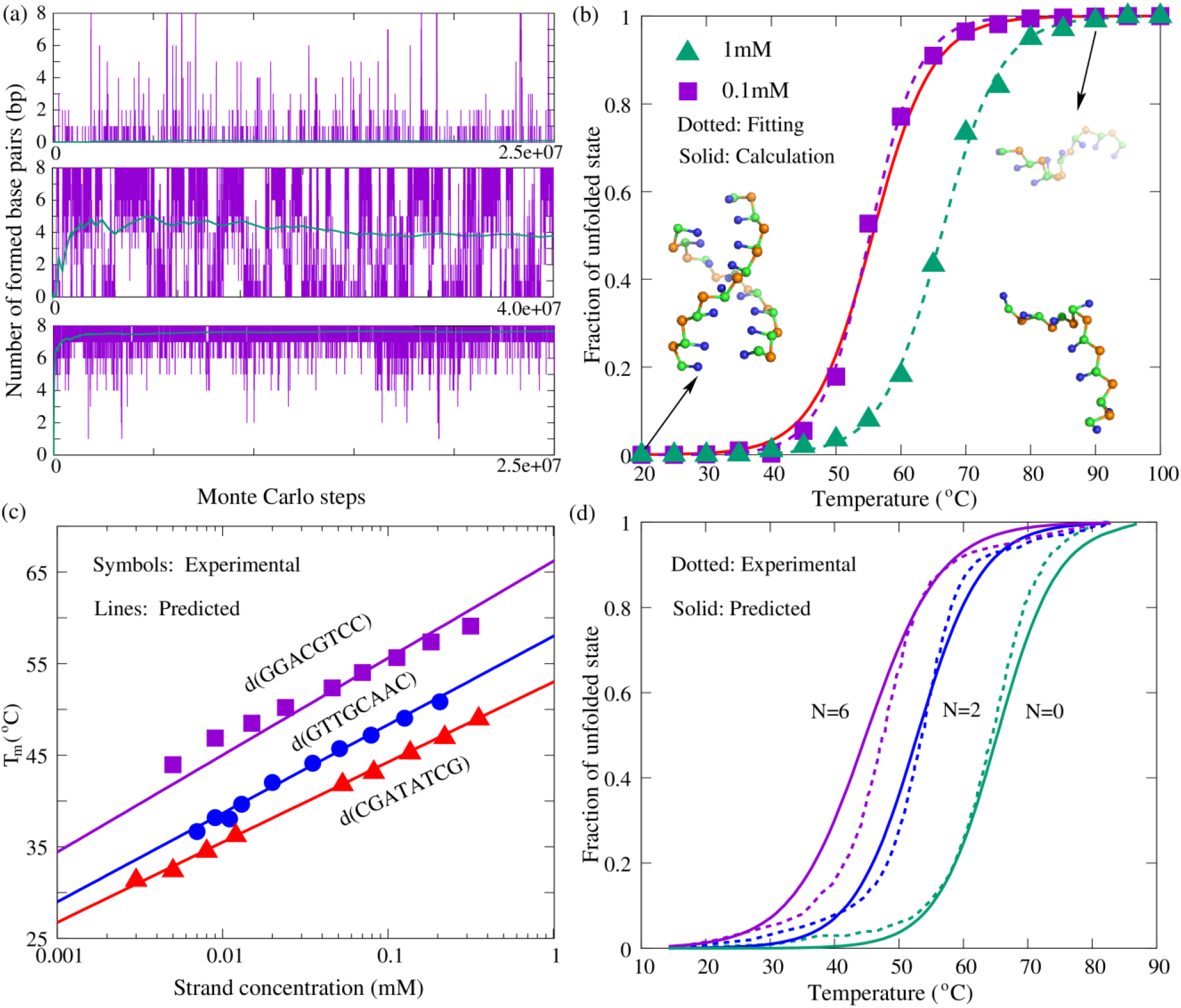
The stability predictions for dsDNAs in the present model. (a) The time-evolution of the number of base-pairs for a dsDNA (sequence: (GGACGTCC)_2_; strand concentration: 1mM) at different temperatures (90°C, 65°C, 40°C from top to bottom, respectively) in 1M NaCl solution. (b) The fractions of unfolded state *f* as functions of temperature for the dsDNA in (a). Green triangle: predictions at high strand concentration (1mM). Purple square: predictions at experimental strand concentration (0.1mM). Two dotted lines are the fitted melting curves to the corresponding predicted data. The solid line is calculated through Eq. 7. Ball-stick: the typical 3D structures predicted at low and high temperatures shown with PyMol (http://www.pymol.org). (c) The melting temperatures (*T*_m_’s) as functions of strand concentration for three dsDNAs: purple, (GGACGTCC)_2_, blue, (GTTGCAAC)_2_, and red, (CGATATCG)_2_ at 1M [Na^+^]. Symbols: experimental results (70). Lines: predictions from the present model. (d) The predicted (solid lines) and experimental (dotted lines) (86) melting curves for the dsDNA harboring symmetric internal loops with sequences of CTCGTC(T)_N_CAGTGC/GCACTG(T)_N_GACGAG in 1M NaCl solution. Green: N=0, i.e., the double helix without internal loop. Blue: N=2. Purple: N=6.

To examine the sequence effect, 27 dsDNAs (8-36nt) with different sequences have been studied with the present model. The sequences, strand concentrations, and the predicted/experimental melting temperatures are listed in Table 1. Here, all dsDNAs are assumed at 1M [Na^+^] to make comparisons with corresponding experimental data. As shown in Table 1, the *T*_m_ values of extensive dsDNAs from the present model are very close to the experimental measurements with a mean deviation of 1.5°C and maximal deviations < 3.0°C, which indicates that the present model with the sequence-dependent base-stacking/pairing potential can make successful predictions on the stability for dsDNA of extensive sequences/lengths. Furthermore, due to the involvement of coaxial stacking potential, the present model can also provide reliable stability for dsDNA with bulge/internal loops, For example, for 4 dsDNAs with bulge loops and 5 dsDNAs with internal loops, the mean deviation of predicted *T*_m_’s from the experiments is only 1.8°C; see Table 1, and the predicted melting curves for the dsDNAs with internal loops of different lengths are just slightly broader than the experiments (86) (Fig. 4d).

#### For ssDNA with various lengths/sequences

Beyond the dsDNA, the stability of ssDNA can also be captured by the present model. As shown in Fig. 5, for DNA hairpins (GCGC(T)_N_GCGC) with different loop lengths (*N*=3-9), the predicted thermal unfolding curves at 0.1M [Na^+^] agree reasonably with the experiments, despite that the predicted *T*_m_ (~78°C) for the hairpin with a small loop (e.g., *N*=3) is rather lower than the experimental value (~80.7°C), while it is a little higher (~58.1°C *vs* ~56.2°C) for large hairpin loops (e.g., *N*=9) (89). Moreover, 24 ssDNAs including pseudoknot are used to verify the ability of the present model for sequence effect on stability; see Table 2. In order to compare with experiments (36,87–93), all these predictions are at corresponding experimental ion conditions. As shown in Table 2, for 24 ssDNAs with different sequences and lengths (11-34nt), the mean/maximal deviation of *T*_m_ between predicted and experimental is ~2.1°C/~3.8°C, which suggests that the effect of sequence on ssDNA stability can also be well described by the present model. It is worth noting that due to the lack of stacking interactions between unpaired bases, the present model cannot distinguish the stability of DNAs with same stems but different loop sequences (e.g., GCGC(T)_5_GCGC *vs* GCGC(A)_5_GCGC), and yet the stability of them generally differ somewhat (9,88,92).

**Figure 5.**
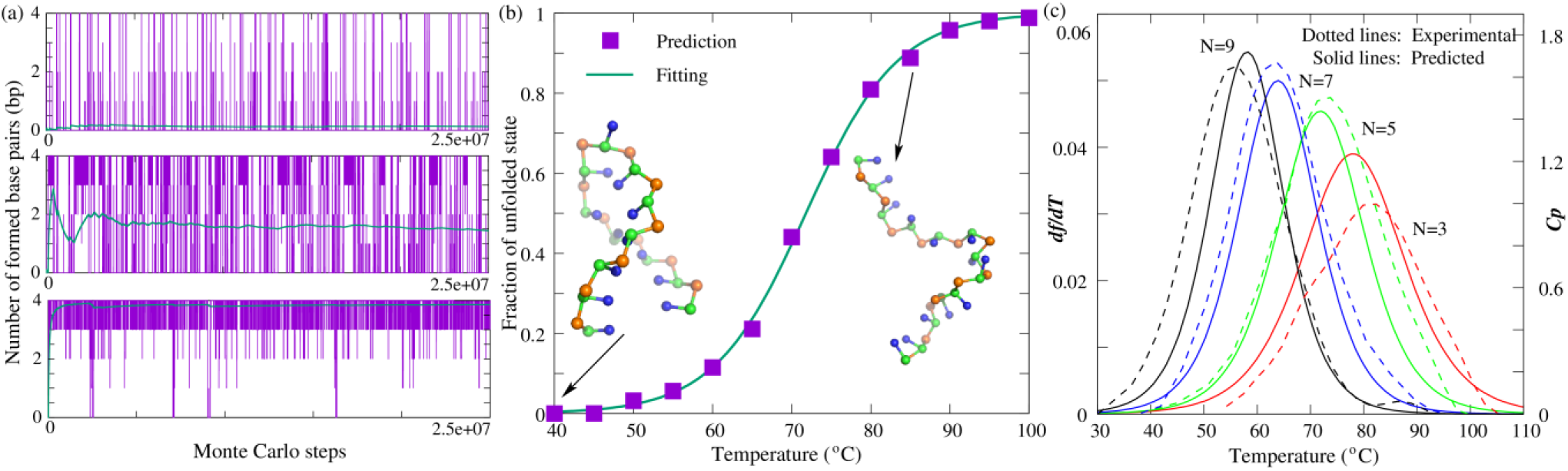
The stability predictions for ssDNA hairpins in the present model. (a) The time-evolution of the number of base-pairs for a simple hairpin (GCGC(T)_5_GCGC) at different temperatures (90°C, 70°C, 50°C from top to bottom, respectively) in 0.1M NaCl solution. (b) The fraction of unfolded state *f* as a function of temperature for the hairpin in (a). Symbols: predictions from the present model. Line: fitted melting curve to the predicted data through Eqs. 4–6. Ball-stick: the typical 3D structures predicted at low and high temperatures shown with PyMol (http://www.pymol.org). (c) The comparisons between predictions (solid lines) and experiments (dotted lines) for four DNA hairpins (GCGC(T)_N_GCGC) with different loop lengths at 0.1M [Na^+^]. Red: N=3. Green: N=5. Blue: N=7. Black: N=9. *df*/*dT*: the first derivative of predicted *f* with the temperature. *C*_*p*_: the heat capacity from experiment (87).

Specifically, we made additional predictions for the stability of two more complex ssDNAs: a pseudoknot and a chain with two hairpins at two ends; see Fig. 6a. As shown in Table 2 and Fig. 6, for the ssDNA with two hairpins, two melting temperatures (*T*_m1_ and *T*_m2_) of the corresponding transitions are successfully predicted by the present model, with the deviations of ~2.1°C and ~1.1°C from experimental data, respectively. Since the hairpin at 3’ end contains fewer G-C pairs than the other (Fig. 6a), it melts at a significantly low temperature in comparison to the 5’ end hairpin (89). For the DNA pseudoknot at 0.1M [Na^+^], the predicted *T*_m1_ and *T*_m2_ are ~48.8°C and ~72.0°C, respectively, which also agree well with the experimental data (~52.6°C and ~70.7°C) (87); see Table 2, and the comparison between predicted and experimental thermal unfolding curves can be found in Fig. 6c. In the predicted curve, the first transition that is from folded pseudoknot state to intermediate hairpin state is more significant than that form experiment. One possible reason is that noncanonical interactions such as triple base interactions between loops and stems and self-stacking in loop nucleotides, which are common in RNA/DNA pseudoknots (68,87), are neglected by the present model, leading to a relatively simple unfolding energy surface. Even so, the comparison with the experiment still suggests that the present model can be reliable in predicting thermal stability for DNA pseudoknots in monovalent ion solutions, and it is noted that the present model can also provide 3D structures for the pseudoknot at different temperatures from the sequences.

**Figure 6.**
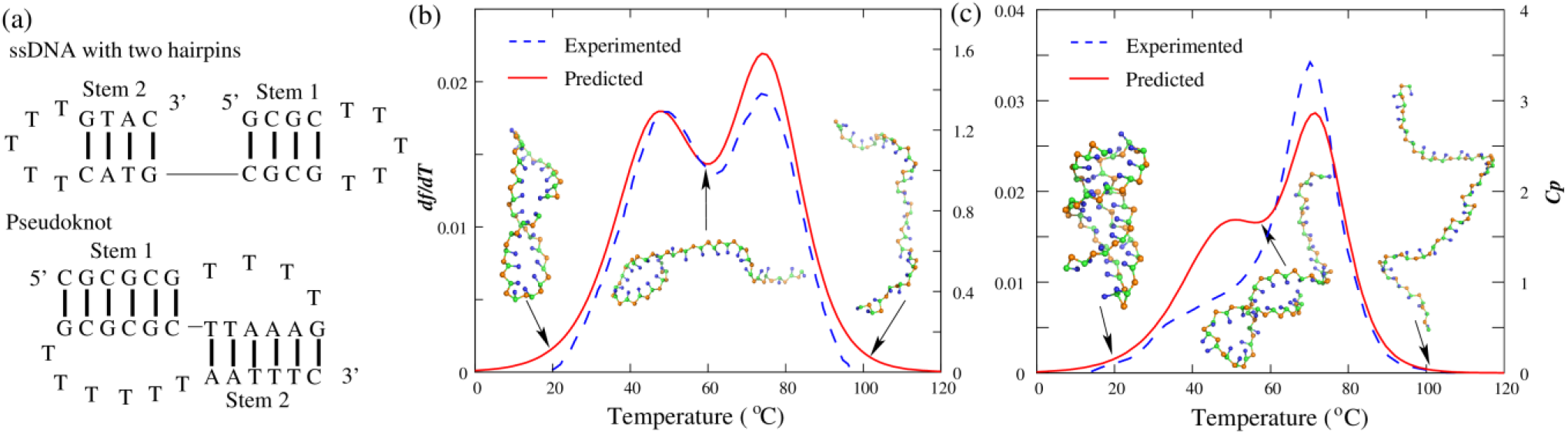
The stability prediction for an ssDNA with two hairpins and a DNA pseudoknot in the present model. (a) The schematics of secondary structure for the ssDNA with two hairpins (top) and the DNA pseudoknot (bottom). (b, c) The comparisons between predictions (solid lines) and experiments (dotted lines) for the two ssDNAs in (a). *df*/*dT*: the first derivative of predicted *f* with the temperature. *C*_*p*_: the heat capacity from experiments (87,89). Ball-stick: the typical 3D structures predicted at different temperatures shown with PyMol (http://www.pymol.org).

### Monovalent/divalent ion effects on stability of dsDNA/ssDNA

Due to the high density of negative charges on the backbone, DNA stability is sensitive to the ionic condition of the solution, while the effect of ions, especially divalent ions (e.g., Mg^2+^), is generally ignored in the existing DNA CG models (43–50). Here, we employed the present model to examine the monovalent/divalent ion effects on the thermal stability of dsDNA and ssDNA.

#### Monovalent ion effect

For each of the three dsDNAs with different lengths (6bp, 10bp, and 15bp), we performed simulations over a broad range of monovalent ion concentrations ([Na^+^]: 0.01M-1.0M), and calculated the melting temperatures at different [Na^+^]’s. As shown in Fig. 7a, the increase of [Na^+^] enhances the dsDNA folding stability due to the stronger ion neutralization (61,62), and the predicted melting temperatures for the three dsDNAs are well in accordance with the experiment results (84,94), with the mean deviation of ~1.4°C. Fig. 7a also shows that the [Na^+^]-dependence of *T*_m_ is stronger for longer dsDNA, which could be caused by the larger buildup of negative charges during base pair formation of longer dsDNA (62,73).

**Figure 7.**
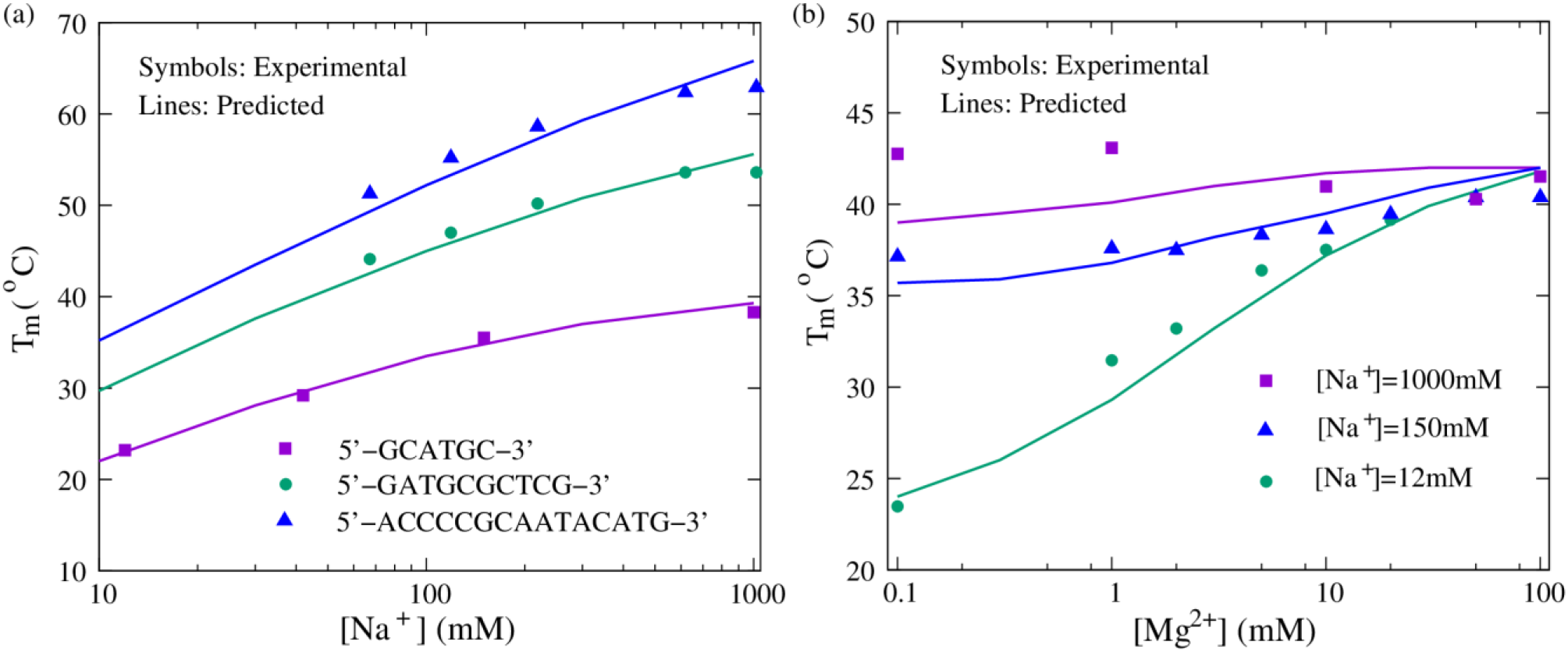
The comparisons of stability between predictions (lines) and experiments (symbols) for dsDNAs in monovalent/divalent ion solutions. (a) The melting temperatures (*T*_m_’s) as functions of [Na^+^] for three dsDNAs with sequences of (GCATGC)_2_, (GATGCGCTCG)_2_, (ACCCCGCAATACATG)_2_ from bottom to top, respectively. Symbols: experimental data (84,94). Lines: predictions from the present model. (b) The melting temperatures (*T*_m_’s) as functions of [Mg^2+^] for the dsDNA with sequence of (GCATGC)_2_ at different [Na^+^]’s: 0.012M, 0.15M, and 1M from bottom to top, respectively. Symbols: experimental data (84). Lines: predictions from the present model.

Although Table 2 has indicated that the present model can make reliable predictions for ssDNA stability at various [Na^+^]’s, we further used a simple DNA hairpin (GCGC(T)_N_GCGC) with different loop lengths (N=5, 7, and 9) to test monovalent ion effect on stability in the present model. As shown in Fig. 8a, for the hairpin with small loops (e.g., N=5 and 7), the difference of predicted *T*_m_ from the experiments over a wide range of [Na^+^]’s is very small (e.g., mean/maximal deviation of ~1.5°C/~1.0°C), and for the loop length of 9, our predictions are slightly larger than the experimental data only at high [Na^+^]’s; e.g., ~4.0°C higher at ~0.1M [Na^+^] (87). The results on the stability of ssDNA and dsDNA in monovalent ion solutions reveal that it is a very effective way of involving the electrostatic interaction for DNA in the present model through the combination of the Debye-Huckel approximation and the concept of counterion condensation, which has also been validated by the TIS model (50,51).

**Figure 8.**
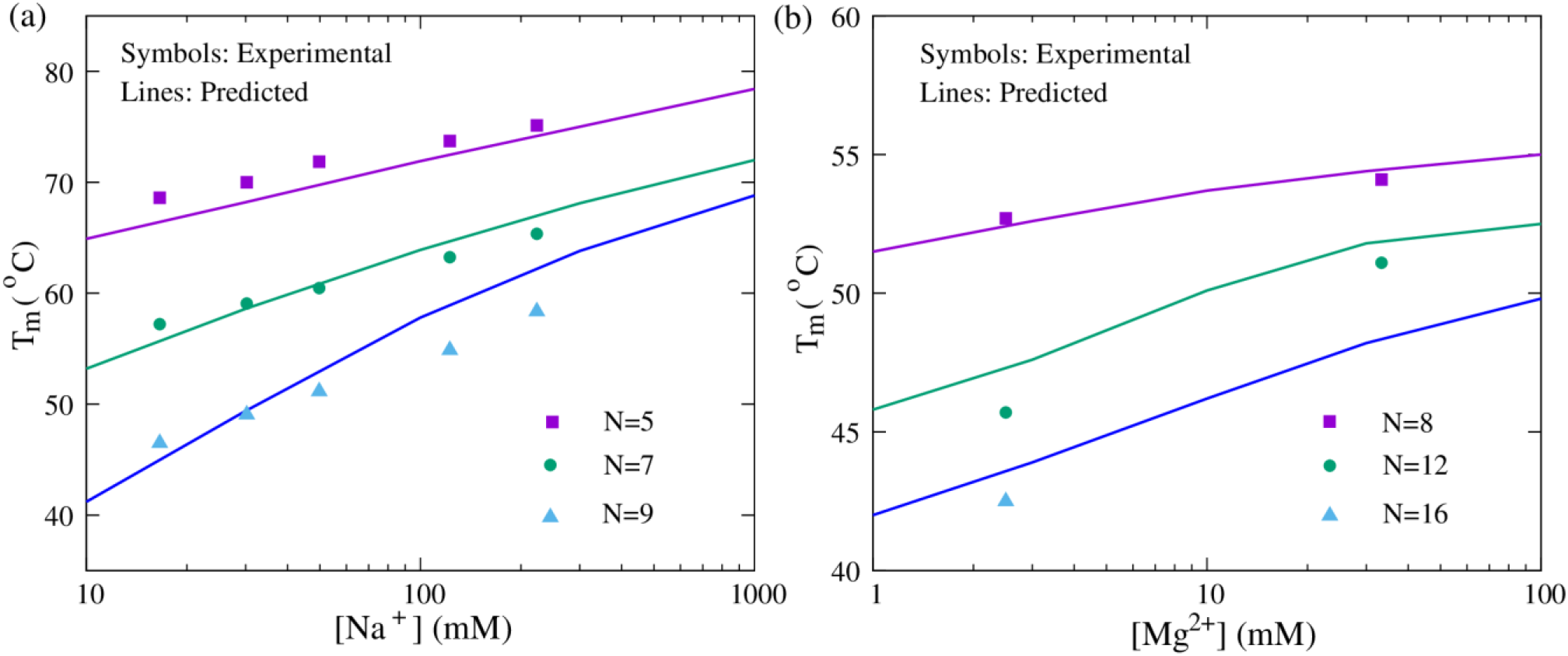
The comparisons of stability between predictions (lines) and experiments (symbols) for ssDNAs in monovalent/divalent ion solutions. (a) The melting temperatures (*T*_m_’s) as functions of [Na^+^] for the DNA hairpins (GCGC(T)_N_GCGC) with different loop lengths: 5, 7, and 9 from top to bottom, respectively. Symbols: experimental data (87). Lines: predictions from the present model. (b) The melting temperatures (*T*_m_’s) as functions of [Mg^2+^] for the DNA hairpins (CGGATAA(T)_N_TTATCCG) with different loop lengths: 8, 12, and 16 from top to bottom, respectively. Symbols: experimental data (93). Lines: predictions from the present model.

#### Divalent ion effect

Remarkably, one important feature of the present model is that combining the counterion condensation theory and the results from the TBI model; see Eq. 3, it can also be used to simulate DNA folding in mixed monovalent/divalent ion solutions. For one dsDNA with the sequence of (GCATGC)_2_ and one ssDNA hairpin with various lengths of the loop (CGGATAA(T)_N_TTATCCG), we made massive predictions in mixed Na^+^/Mg^2+^ solutions and compared the melting temperatures with the corresponding experimental results (84,93); see Figs. 7b and 8b. The comparisons of *T*_m_’s over a wide range of [Mg^2+^] are in line with the experiments, whether for the dsDNA at different [Na^+^]’s (i.e., 0.012M, 0.15M, and 1M) or for the ssDNA with different lengths of the loop, which suggests that the present model can nearly make quantitative predictions for the stability of DNAs in mixed ion solutions from their sequences, even though the ion effect is involved implicitly.

Furthermore, the competition between Na^+^ and Mg^2+^ on DNA stability can also be captured by the present model. For example, for dsDNA at 0.012M [Na^+^] (Fig. 7b), when [Mg^2+^] is very low (e.g., <0.3mM), Na^+^ dominates the stability of the dsDNA, while the increase of [Mg^2+^] enhances the stability significantly. This is because the bindings of Na^+^ and Mg^2+^ are generally anti-cooperative and Mg^2+^-binding is more efficient in stabilizing DNA structures (62,72). Naturally, as [Na^+^] increases, the negative charge on DNA is strongly neutralized, and consequently, the effect of Mg^2+^ appears weak.

## Discussion

In this work, we have proposed a novel three-bead CG model to predict 3D structure and stability for both ssDNA and dsDNA in ion solutions only from the sequence. As compared with the extensive experiments, we have demonstrated that, (1) The present model can successfully predict the native-like 3D structures for ssDNAs and dsDNAs with an overall mean (minimum) RMSD of ~3.4Å (~1.9Å) from corresponding experimental structures, and the overall prediction accuracy of the present model is slightly higher than the existing models; (2) The present model can make reliable predictions on stability for dsDNAs with/without bulge loops and ssDNAs including pseudoknots, and for 51 DNAs with various lengths and sequences, the predicted melting temperatures are in good accordance with extensive experiment data (i.e., mean deviation of ~2.0°C); (3) The present model with implicit electrostatic potential can also reproduce the stability for ssDNAs/dsDNAs at extensive monovalent or mixed monovalent/divalent ion conditions, with the predicted melting temperatures consistent with the available experiments.

Nonetheless, the present model has several limitations that should be overcome in future model development. For example, the present model failed to predict native-like structures for more complex DNAs such as that with three-way junction and cannot distinguish the stability for DNAs with different loop sequences, which suggest that possible noncanonical interactions (e.g., base triple interactions between loops and stems, self-stacking in loop nucleotides and special hydrogen bonds involving phosphates and sugars should be further taken into account (2,50,82). Furthermore, a more efficient sampling algorithm such as replica-exchanged MC or MC with umbrella sampling, as well as suitable structure constraints should be introduced to the model assembly for large DNAs (e.g., nano-architectures) (46–49,50), and accordingly, an accurate score function like statistical potential used for RNA and protein could be required to evaluate predicted DNA candidate structures (95–100). In addition, the 3D structure predicted by the present model is at the CG level, and it is still necessary to reconstruct all-atomistic structures based on the CG structures for further practical applications. After these further developments, a user-friendly web server would be further freely available, allowing users to predict 3D structure and stability for DNAs in ion solutions from sequence or given constraints.

## Data Availability Statement

All relevant computational data are within the paper and its Supporting information files. Experimental data are publicly available from published papers cited in the work. The 3dRNA/DNA web server used in this work is available by http://biophy.hust.edu.cn/new/3dRNA/create.

## Supporting information

**S1 Text**. The force field of the present model, the melting temperature calculation for dsDNAs at low strand concentrations, and the additional figures.

## Acknowledgments

We are grateful to Profs. Zhi-Jie Tan (Wuhan University), Wenbing Zhang (Wuhan University), and Yaoqi Zhou (Shenzhen Bay Laboratory) for valuable discussions, and we would like to acknowledge computing resources from the Shenzhen Bay Laboratory Supercomputing Center.

## Author Contributions

Conceptualization: Ya-Zhou Shi, Zi-Chun Mu, Ya-Lan Tan.

Data curation: Zi-Chun Mu, Ya-Lan Tan.

Formal analysis: Ya-Zhou Shi, Zi-Chun Mu.

Funding acquisition: Ya-Zhou Shi, Ben-Gong Zhang.

Investigation: Zi-Chun Mu, Ya-Zhou Shi, Jie Liu.

Methodology: Ya-Zhou Shi, Zi-Chun Mu.

Supervision: Ya-Zhou Shi.

Validation: Zi-Chun Mu, Ya-Zhou Shi, Ya-Lan Tan.

Writing – original draft: Zi-Chun Mu, Ya-Lan Tan, Ya-Zhou Shi.

Writing – review & editing: Zi-Chun Mu, Ya-Zhou Shi, Jie Liu.

